# Platinum nanoparticles inhibit intracellular ROS generation and protect against Cold Atmospheric Plasma-induced cytotoxicity

**DOI:** 10.1101/2021.02.18.431888

**Authors:** Sebnem Gunes, Zhonglei He, Renee Malone, Patrick J Cullen, James F Curtin

**Affiliations:** BioPlasma Research Group, School of Food Science and Environmental Health, Technological University Dublin, Dublin, Ireland; Nanolab, FOCAS Research Institute, Technological University Dublin, Dublin, Ireland; Environmental, Sustainability and Health Research Institutes, Technological University Dublin, Dublin, Ireland; School of Chemical and Biomolecular Engineering, University of Sydney, Australia

**Keywords:** Cold Atmospheric Plasma, Free Radicals, Platinum Nanoparticles, Antioxidants, Cancer Treatment, Nanotherapy

## Abstract

Platinum nanoparticles (PtNPs) have been investigated for their antioxidant abilities in a range of biological and other applications. The ability to reduce off-target CAP cytotoxicity would be useful in Plasma Medicine, however, little has been published to date about the ability of PtNPs to reduce or inhibit the effects of CAP. Here we investigate whether PtNPs can protect against CAP-induced cytotoxicity in cancerous and non-cancerous cell lines. PtNPs were shown to dramatically reduce intracellular reactive species (RONS) production in human U-251 MG cells. However, RONS generation was unaffected by PtNPs in medium without cells. PtNPs protect against CAP induced mitochondrial membrane depolarization, but not cell membrane permeabilization which is a CAP-induced RONS-independent event. PtNPs act as potent intracellular scavengers of reactive species and can protect both cancerous U-251 MG cells and non-cancerous HEK293 cells against CAP induced cytotoxicity. PtNPs may be useful as a catalytic antioxidant for healthy tissue and for protecting against CAP-induced tissue damage.

**Graphical Abstract:** PtNPs are potent catalase and superoxide dismutase mimetics which makes them strong antioxidant candidates for the protection of cells against oxidative stress. CAP was generated using a Dielectric Barrier Device (DBD) system with a voltage output of 75 kV at a frequency of 50 Hz. A range of concentrations of 3nm uncoated PtNPs combined with CAP were examined in human U-251 MG Glioblastoma (GBM) cells and non-cancerous human embryonic kidney HEK293 cells. The protective effects of PtNPs against CAP were explored using several biochemical indicators of oxidative stress and cytotoxicity.

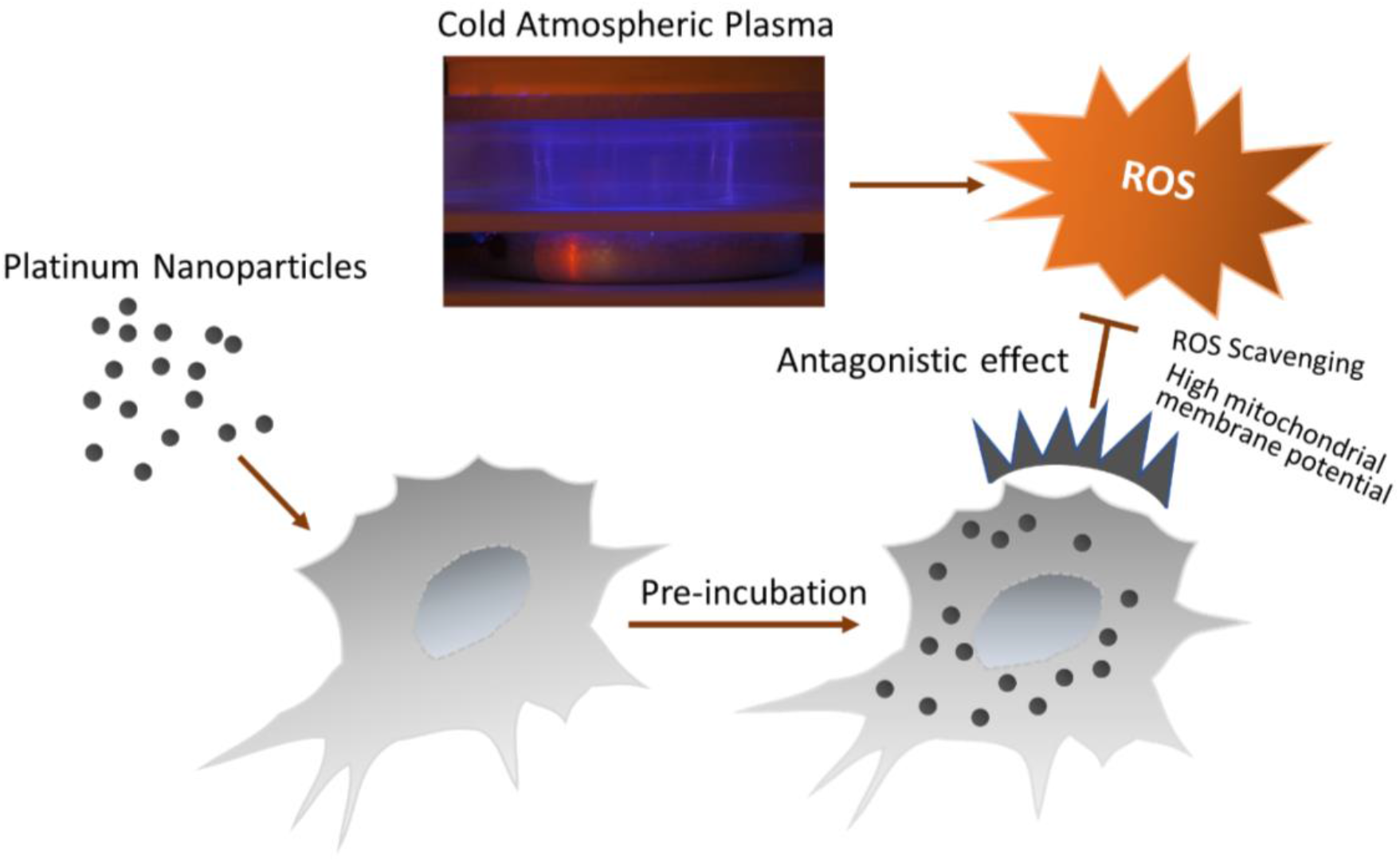

## Background

Cold atmospheric plasma (CAP), also known as non-thermal atmospheric plasma (NTAP), has emerged as a promising technique for biomedical applications, including sterilization, wound healing and cancer therapy^1,2^. CAP is known to generate bioactive short- and long-lived reactive oxygen species (ROS) and reactive nitrogen species (RNS), as well as a unique physical environment, including heat generation, pressure gradients, electrostatic and electromagnetic fields^3,4^. In many types of tumours, cancer cells have been found to possess a higher intracellular ROS level compared to normal cells, due to their more rapid proliferation and higher, altered metabolic activities^5^. Therefore, the antioxidant system in cancer cells is under a heavier workload, potentially causing a higher sensitivity of cancer cells to oxidative stress compared to normal cells, which can be targeted for the development of specific therapies. For example, several anticancer drugs, such as imexone^6^, motexafin gadolinium^7^ and nitric oxide-donating aspirin^8^, have been developed to increase ROS level in cancer cells beyond their tolerable allowance to trigger cell death. CAP has been demonstrated to have selectively anti-tumour effects against many cancer cell lines^9,10^. However, even with non-immediate damage to healthy tissues by precise CAP treatment, the increase of the ROS levels in surrounding tissues may induce DNA damage and cause secondary malignancies, which can be a potential side effect induced by CAP^10,11^.

Recently, it has been demonstrated that CAP treatment has synergistic anti-cancer effects in combination with various nanoparticles, including gold nanoparticles^12^, silver nanoparticles^13^, iron nanoparticles^14^ and iron oxide-based magnetic nanoparticles^15^.

Antioxidants protect against cancer occurrence and cardiovascular diseases^16,17^ by scavenging ROS. However, antioxidants as a redox agent may also promote free radicals and secondary ROS and the excess amount of these increases the cancer risk^18,19^. Unlike platinum-based compounds, such as cisplatin that are cytotoxic agents^20,21^, PtNPs display promising antioxidant activity, functioning as superoxide dismutase (SOD)/catalase mimetics^22,23^ and catalyse the decomposition of H_2_O_2_^24,25^ and ROS^26^. H_2_O_2_ is a long-lived reactive species. In cells it can interact with DNA and other biomolecules and can cause biomacromolecular damage^27,28^. Therefore, controlling and removal of excess H_2_O_2_ in neighbouring healthy tissues will improve the selectivity of CAP as a treatment for cancer.

Many previous *in vivo* and *in vitro* studies proposed that coated or uncoated PtNPs have negligible cytotoxic effects with cell compatible materials^29–31^. PtNPs are capable of scavenging ROS and show SOD catalytic activity^26,32,33^. PtNPs scavenge O_2_^-^ (superoxide anion radicals) and .OH (hydroxyl radicals) acting as antioxidants^30,33^. It has been reported that PtNPs reduce oxidative stress-mediated cell damage and cell death^34,35^. Therefore, they may act as an antioxidant against CAP treatment. However, due to the wide range of CAP devices and types, the variety of cell types, different sizes, shapes, purity, and coating of PtNP and variability of experimental conditions, the detailed mechanism needs further research.

In this study, we determined the effects of PtNPs combined with CAP in U-251 MG and HEK293 cells. We demonstrate that PtNPs act as intracellular ROS generation scavengers and protect against CAP-induced cytotoxicity. PtNPs may be useful as a catalytic antioxidant for healthy tissue and protect against CAP-induced tissue damage. Our results indicate that PtNP could be developed as a safe method to reduce the side effects and potential risks of CAP and other ROS-related therapies on adjacent healthy tissues during treatment.

## Methods

### Cell Culture

The human brain glioblastoma cancer cell line (U-251 MG) was obtained from Dr Michael Carty (Trinity College Dublin) and the human embryonic kidney cell line (HEK293) from Dr Darren Fayne (Trinity College Dublin). Cells were cultured in Dulbecco’s Modified Eagle’s Medium-high glucose (Merck) supplemented with 10% fetal bovine serum (Merck) and 1% penicillin (Thermo Fisher Scientific) and maintained in a humidified incubator containing 5% (v/v) CO_2_ atmosphere at 37°C in TC flask T25, standard for adherent cells (Sarstedt). Culture medium was changed every 2-3 days upon reaching around 80% confluence. Cells were routinely sub-cultured in new flasks using a 0.25% Trypsin-EDTA solution (Merck). Platinum, nanoparticle dispersion, 3 nm, was purchased from Merck.

### Alamar blue cell viability assay

Cell viability was analysed using the Alamar blue assay (Thermo Fisher Scientific). U-251 MG and HEK293 cells were seeded at a density of 1×10^4^ cells/well (100 μl culture medium per well) into 96-well plates (Sarstedt) and were incubated overnight. Medium was removed and fresh culture medium containing 0– 100 μg/ml PtNPs was added and incubated overnight. Medium was removed for DBD (DIT-120 plasma device) CAP (75 kV, 50 s) treatment at 70–80% confluences and fresh culture medium was replaced immediately following CAP treatment. Following a 24 h incubation at 37 °C, the cells were rinsed once with phosphate buffered saline (Merck), incubated for 3 h with 10% Alamar blue in culture medium solution. As a second method, we treated U-251 MG cells with PtNPs immediately prior to CAP treatment. An Alamar Blue cell viability assay was performed 48 hours after CAP treatment. Fluorescence was measured (excitation, 530 nm; emission, 595 nm) by a Victor 3 V 1420 microplate reader (Perkin Elmer).

### Measurement of ROS induced by CAP

Reactive oxygen species generation induced by CAP were detected using a cell permeable oxidant sensitive fluorescent dye 2,7-dichlorodihydrofluorescein diacetate (H_2_DCFDA) (Thermo Fisher Scientific). U-251 MG cells were seeded in 35 × 10 mm Petri dishes (Sarstedt) at a density of 1×10^5^ cells/ml. After 24h, growth medium was removed and PtNPs (5 μg/ml and 0.032 μg/ml) in medium were added and then incubated overnight. Culture medium was replaced with fresh serum-free medium containing 25 μM H_2_DCFDA and cells were incubated for 30 min at 37 °C. Cells were washed with fresh medium once and then with PBS twice, and then treated with CAP at 75 kV for 50 s. Following CAP treatment, cells were incubated with fresh medium for 10 minutes at 37 °C before cell harvesting. To prepare aliquots, all floating and attached cells were collected by trypsinisation. All liquids, including medium, washing PBS and trypsin-cell suspension, were transferred into one tube and centrifuged at 1200 rpm for 5 min. Cells were resuspended in PBS and BD Accuri™ C6 Plus flow cytometry (BD Bioscience) was used to detect and measure fluorescence. Flow analysis was carried out with a 488 nm laser for excitation and FL1 standard filter for H_2_DCFDA measurement.

### Detection of cell membrane damage

Viable and dead cells were recorded using Propidium Iodide (PI) staining (Merck). U-251 MG cells were plated in 35 × 10mm Petri dishes (Sarstedt) at a density of 1×10^5^ cells/ml. After 24h, growth medium was removed and PtNPs (5 μg/ml and 0.032 μg/ml) in medium were added and incubated for a further 24 h. Medium was removed and cells were treated directly to CAP 75kV for 50s and, 24h after CAP treatment, cells were collected by trypsinization, and pelleted by centrifugation at 1200 rpm for 5 minutes. The pellet was resuspended in 0.5 ml PBS and cells were stained with 0.4 μg/ml PI for 5 minutes. The fluorescence of PI was then measured using BD Accuri™ C6 Plus flow cytometry at FL2 (585/40 nm) standard filter.

### Measurement of mitochondrial membrane potential

Dual-emission potential-sensitive fluorescence dye JC-1 (Merck) was used to measure mitochondrial membrane potential of cells following CAP treatment. U-251 MG cells were seeded in 35 × 10 mm Petri dishes (Sarstedt) at a density of 1×10^5^ cells/ml. After 24h, growth medium was replaced with PtNPs (5 μg/ml and 0.032 μg/ml) in medium and incubated for a further 24 h. Medium was removed, and the cells were exposed to CAP directly at 75 kV for 50 s. Following CAP treatment, cells were incubated with fresh medium overnight at 37 °C. Medium was removed and 2.5 μg/ml JC-1 in medium was added to cells. After 10-minute incubation time at room temperature and absence of light, cells were washed with PBS twice and all cells were collected by trypsinization and centrifugation at 1200 rpm for 5 min. The supernatant was removed, and cells were resuspended in 0.3 ml PBS for flow cytometry (BD Bioscience) analysis. Fluorescence intensity was measured using the FL1 (530 nm) and FL2 (585 nm) channels with emission spectral overlap compensation (7% FL1/FL2 and 13% FL2/ FL1).

### Statistical Analysis

Independent experiments were carried out for each data point in triplicate. Prism 6 (GraphPad Software) was used to carry out curve fit and statistical analysis. Data is shown as the % and error bars of all figures are presented using the standard error of the mean (S.E.M). Two-tailed P values were used and the Alpha for all experiments is 0.05. Data points were verified using one-way ANOVA and Tukey’s multiple comparison post-test to determine the significance, as indicated in figures (*P<0.05, **P<0.01, ***P<0.001, ****P<0.0001).

## Results

### CAP-induced antagonistic cytotoxicity of PtNPs on cancer cells

We wished to determine whether PtNP could have a protective effect against CAP-induced cytotoxicity. Dose response curves were established using a range of concentrations (Figure 1A, B) to determine the appropriate nontoxic concentrations of PtNPs in cells with the combination of CAP. Control cells were treated with a range of concentrations of PtNPs in the absence of CAP. An Alamar Blue cell viability assay was performed to determine the extent of cytotoxicity. Approximately 50% loss of cell viability was observed 24 h after U-251 MG cells were exposed to 50 s CAP treatment. As Figure 1A and 1B demonstrate, lower concentrations of PtNPs alone did not cause any toxicity to cells without CAP treatment. Moreover, protection from CAP-induced cytotoxicity was observed when pre-incubated with PtNPs. Compared with CAP only, the significant protective concentrations of PtNP ranged from 4 μg/ml to 0.0512 ng/ml when PtNP were preincubated for 24 h (Figure 1A) and from 4 μg/ml to 0.256 ng/ml when PtNP were added immediately prior to CAP treatment (Figure 1B). As seen from the dose response curves in Supplementary Figure S1a The IC_50_ value of PtNP alone is 8.091 μg/ml (95% confident intervals = 6.408 μg/ml to 10.22 μg/ml). The IC_50_ value of PtNP in combination with 50 s CAP treatment is 8.03 μg/ml (95% confident intervals = 5.663 μg/ml to 11.39 μg/ml). Based on the pre-incubation data, 0.032 μg/ml of PtNPs was selected as a protective concentration and 5 μg/ml of PtNPs to compare with 0.032 μg/ml for the cytotoxicity and protective effects for all future experiments. All cell viability data are normalized relative to control cells.

**Figure 1.**
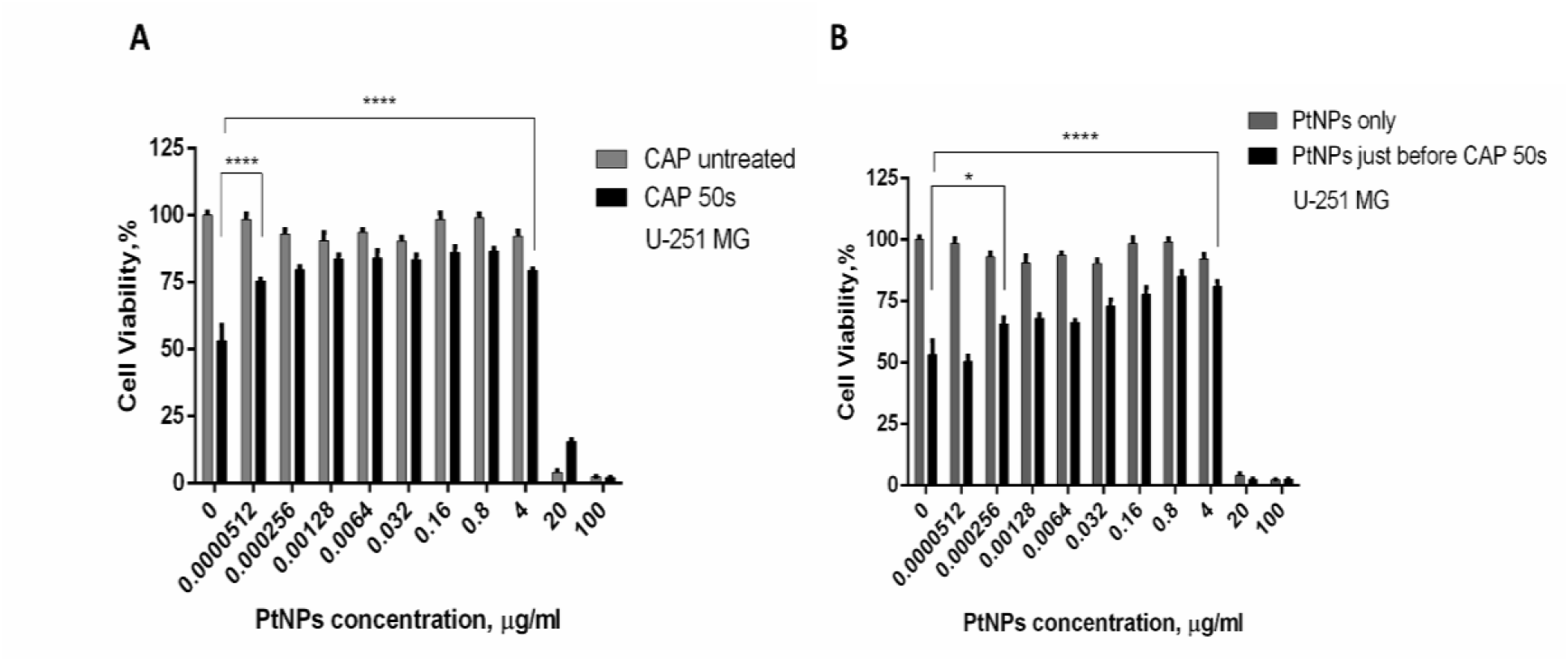
Dose responses for PtNPs treatment. (A) U-251 MG cells were incubated with increasing concentrations (0 ≤ 100 μg/ml) of PtNPs for 24h before CAP treatment and Alamar blue analysis was carried out 24h after CAP treatment (B) U-251 MG cells were treated with corresponding amounts of PtNPs immediately before CAP treatment and Alamar blue cell viability assay was carried out 48h after CAP treatment.

Note that this demonstrates that CAP and PtNPs have an antagonistic cytotoxicity to U-251 MG cancer cell lines.

### Reactive Oxygen Species (ROS) Scavenging Activity of PtNPs in U-251 MG cells

To determine the effects of PtNP on ROS generation by CAP, U-251 MG cells were preloaded with H_2_DCFDA. ROS levels in cells following CAP treatment significantly increased compared to the negative control (Figure 2A) (****P < 0.0001). The levels of intracellular oxidised H_2_DCFDA were decreased when cells were pre-incubated with three different non-toxic concentrations of PtNPs; 0.0512 ng/ml, 0.032 μg/ml, and 5 μg/ml as seen in Figure 2B. The mean fluorescence levels of CAP-treated cells pre-incubated with PtNPs were decreased by a factor of 1.5, 3, and 2.6 times, respectively, compared to CAP-treated cells in the absence of PtNPs (****P < 0.0001).

**Figure 2.**
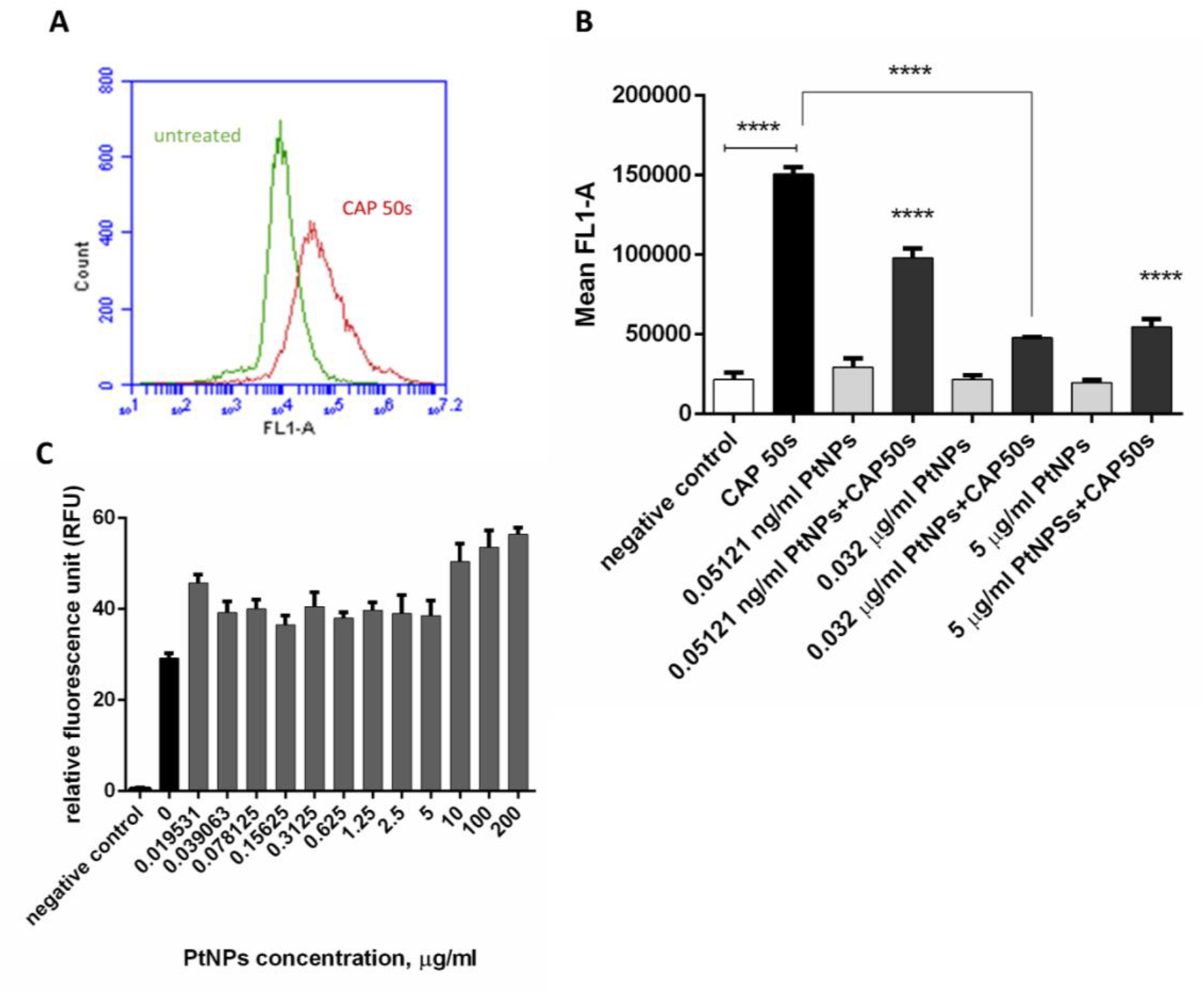
Antioxidant activity. (A) Comparison of ROS production in U-251 MG cells with CAP treatment and untreated control (B) Negative Control (empty column) represents untreated cells, black CAP treatment and others 3 different concentrations of PtNPs (0.05121 ng/ml; 0.032 μg/ml, 5μg/ml) with and without CAP treatment and data was acquired by flow cytometry. (C) Fluorescence level of oxidised H_2_DCFDA was measured in medium without cells after CAP treatment with increasing concentrations (0 ≤ 200 μg/ml) of PtNPs. Data retrieved from the Perkin Elmer microplate reader measures the fluorescence in each well of the microplate and generates readings as arbitrary fluorescent unit values.

In contrast, the presence of PtNPs in culture media only (without cells) was not sufficient to prevent oxidation of H_2_DCFDA in response to 50s CAP treatment suggesting that either localized accumulation and concentration of PtNP, or a role for the intracellular redox machinery, are important for the antioxidant properties displayed (Figure 2C).

### Effect of PtNPs combined with CAP on cell membrane damage

We have previously demonstrated that exposure of U-251 MG cells to CAP DBD induces rapid membrane damage and permeability to dyes such as Propidium Iodide (PI)^36^. To assess the effect of PtNP on membrane damage, U-251 MG cells were stained with PI. PtNPs alone were not observed to cause membrane damage (Figure 3). As expected, a rapid increase in membrane permeability was observed following exposure to CAP. The mean fluorescence in cells with 5 μg/ml PtNPs exposed to the CAP increased by a factor of 1.5, while concentrations of PtNPs 0.032 μg/ml combined by CAP showed no a significant difference, compared to cells following only the CAP treatment. As seen Figure 3A, untreated cells when compared to CAP only and PtNPs combined with CAP showed significant difference (**P < 0.01, ****P < 0.0001). The percentage of viable and dead cells did not show significant difference after PI staining between pre-incubation of PtNPs combined with CAP treatment and CAP 50 s treatment only (Figure 3B). Figure 3C illustrates the emitting fluorescence from viable and dead cells. This observation suggests 50s CAP by the DBD device induces immediate membrane permeabilization accounting for a fraction of the overall cytotoxicity that is not prevented by the presence of PtNPs.

**Figure 3.**
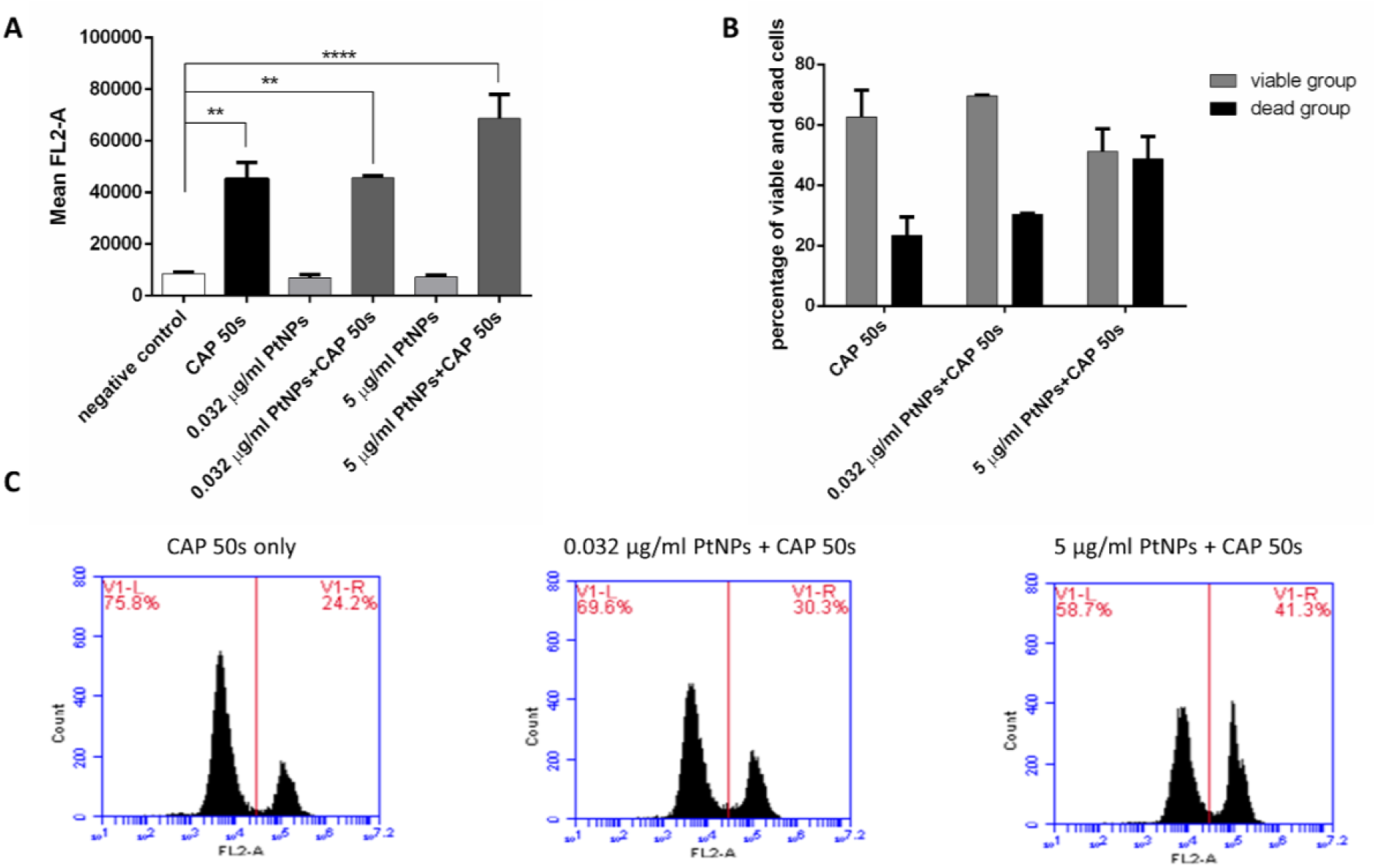
Cell membrane damage. (A) Fluorescence level of PI staining in untreated, CAP-treated, with pre-incubated 0.032 μg/ml and 5 μg/ml of PtNPs alone and the combination of PtNPs and CAP-treated cells. (B) Comparison of the percentage of viable and dead cells with CAP treatment only and 0.032 μg/ml and 5 μg/ml of PtNPs combined with CAP. (C) The percentage of viable, on the left side, and dead, on the right side, cells emitting red fluorescence. Figures represent CAP treatment only, 0.032 μg/ml of PtNPs combined with CAP and 5 μg/ml of PtNPs combined with CAP.

### High mitochondrial membrane potential with PtNPs combined by CAP

Inhibition of ROS generation in cells, but no effect on the immediate membrane damage induced by CAP, suggests that PtNPs play a protective role by modifying the intracellular ROS signaling in cells following exposure to CAP and membrane damage. Mitochondria play an important role in ROS-mediated cell death by both producing and augmenting cellular responses to reactive species. The accumulation of ROS in cells can result in oxidative stress which leads to mitochondrial dysfunction and CAP is known to cause the loss of the mitochondrial membrane potential (ΔΨm)^37^.

We therefore investigated whether PtNP affected mitochondrial membrane depolarization. U-251 MG cells were stained with JC-1. The dimeric form of JC-1 predominates in healthy mitochondria and exhibits a red fluorescence (Figure 4). In contrast, mitochondrial membrane depolarization following CAP treatment led to a shift to green fluorescence indicating that JC-1 was predominantly monomeric. To determine the changes of JC-1 monomer/dimer mean fluorescence ratio in U-251 MG cells, PtNPs concentrations of 0.032 μg/ml and 5 μg/ml with and without 50 s CAP treatment were evaluated. As evident in Figure 4, a low ratio of monomers-to-aggregates was observed in PtNPs treated cells, further indicating that the concentrations of PtNP selected did not have a toxic effect on cells. CAP treatment led to the expected increment of the green/red fluorescence ratio, with a threefold increase in mean fluorescence intensity compared to untreated controls.

**Figure 4.**
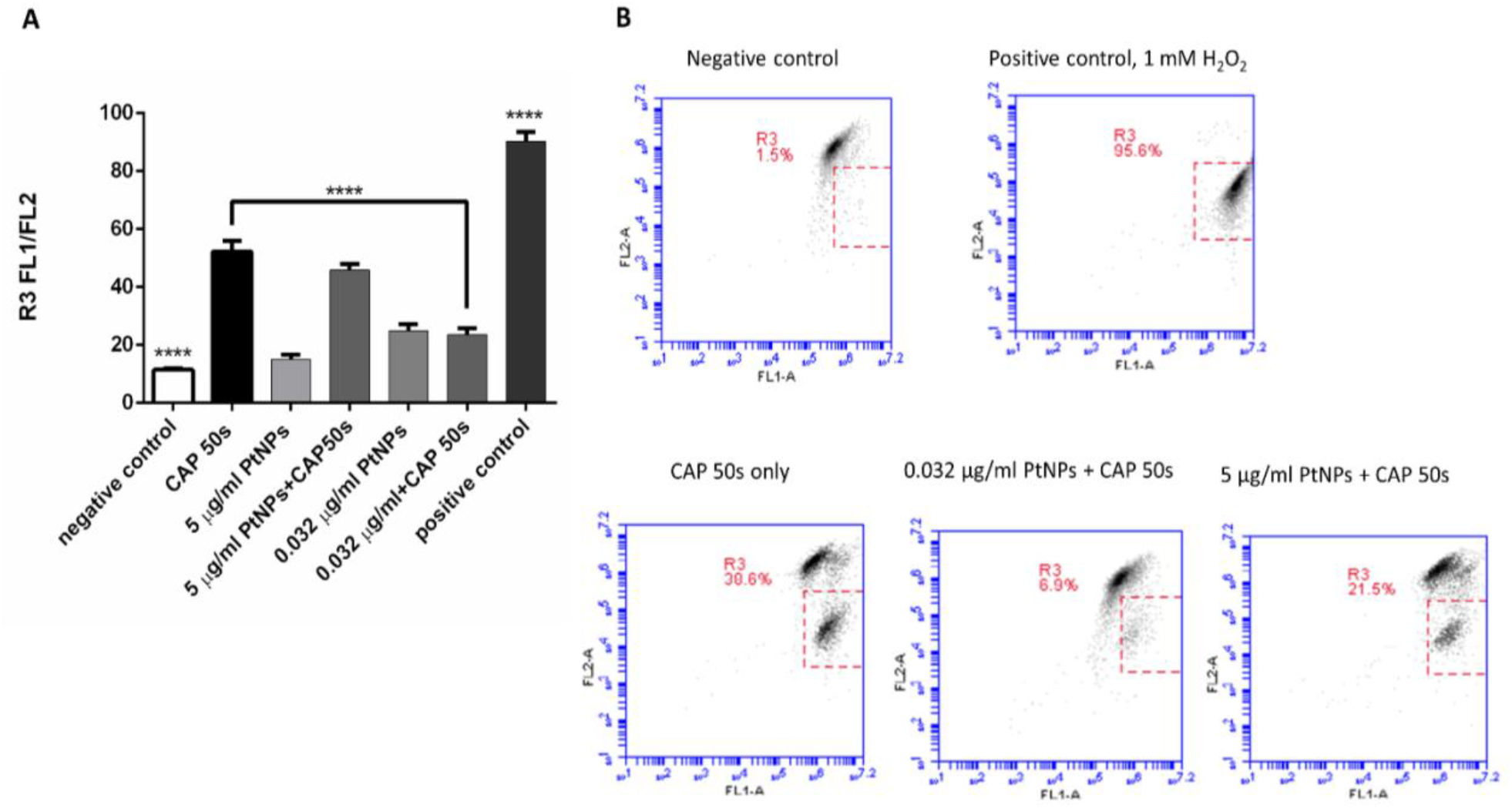
Mitochondrial membrane depolarisation in GBM. (A) Cells emitting green fluorescence and with low mitochondrial membrane potential against cells emitting red fluorescence and with high mitochondrial membrane potential are represented as a bar graph and mean fluorescence ratio of FL1/FL2 was acquired from the gate R3 in panel (B). Bars represent untreated, 0.032 μg/ml and 5 μg/ml of PtNPs combined with and without CAP, CAP treatment alone and 1 mM H_2_O_2_ as positive control groups toxicity comparison. (B) Cells emitting green fluorescence and with low mitochondrial membrane potential are shown in percentage as R3. Figures represent negative control, positive control, CAP treatment only, 0.032 μg/ml of PtNPs combined with CAP and 5 μg/ml of PtNPs combined with CAP.

These results indicate that the ΔΨm is disrupted by CAP treatment and JC-1 remained in its monomeric form. The extent of membrane depolarization by CAP was similar to that observed when cells are exposed to 1mM H_2_O_2_ treatment as a positive control group. Interestingly, the ratio of FL1/FL2 mean fluorescence intensity in cells with PtNPs that are exposed to the CAP treatment for 50 s decreases by a factor of 2.5 and 1.3 with 0.032 μg/ml and 5 μg/ml of PtNPs, respectively, compared to 50 s CAP treatment alone (****P < 0.0001). This observation shows that PtNPs, especially at a concentration of 0.032 μg/ml, support a high retention of mitochondrial membrane potential by removing the disrupting effect of the CAP treatment; therefore, JC-1 moves and aggregates inside the mitochondria.

### CAP-induced antagonistic cytotoxicity of PtNPs on healthy cells

Our results point to the potential of PtNP as a potent inhibitor of cytotoxicity. While this effect would not be desirable in treating cancer cells, the effect would be very beneficial to limit bystander damage if used to protect neighbouring healthy cells and tissues. To investigate whether the effect was also observed in healthy cells, the non-cancerous human embryonic kidney HEK293 cell line was used as contrast to U-251 MG cell line to measure cell viability. We treated non-cancerous cells with a range of concentrations of PtNPs combined with CAP to see effects of PtNPs on healthy cells. Our data indicated that HEK293 cell viability was approximately 60% after 50 sec plasma treatment 24 h post-CAP treatment (Figure 5). Use of PtNPs at concentrations lower than 20 μg/ml did not lead to any measurable cytotoxicity in this cell line, however, at higher concentrations the PtNPs were cytotoxic and decreased cell viability.

**Figure 5.**
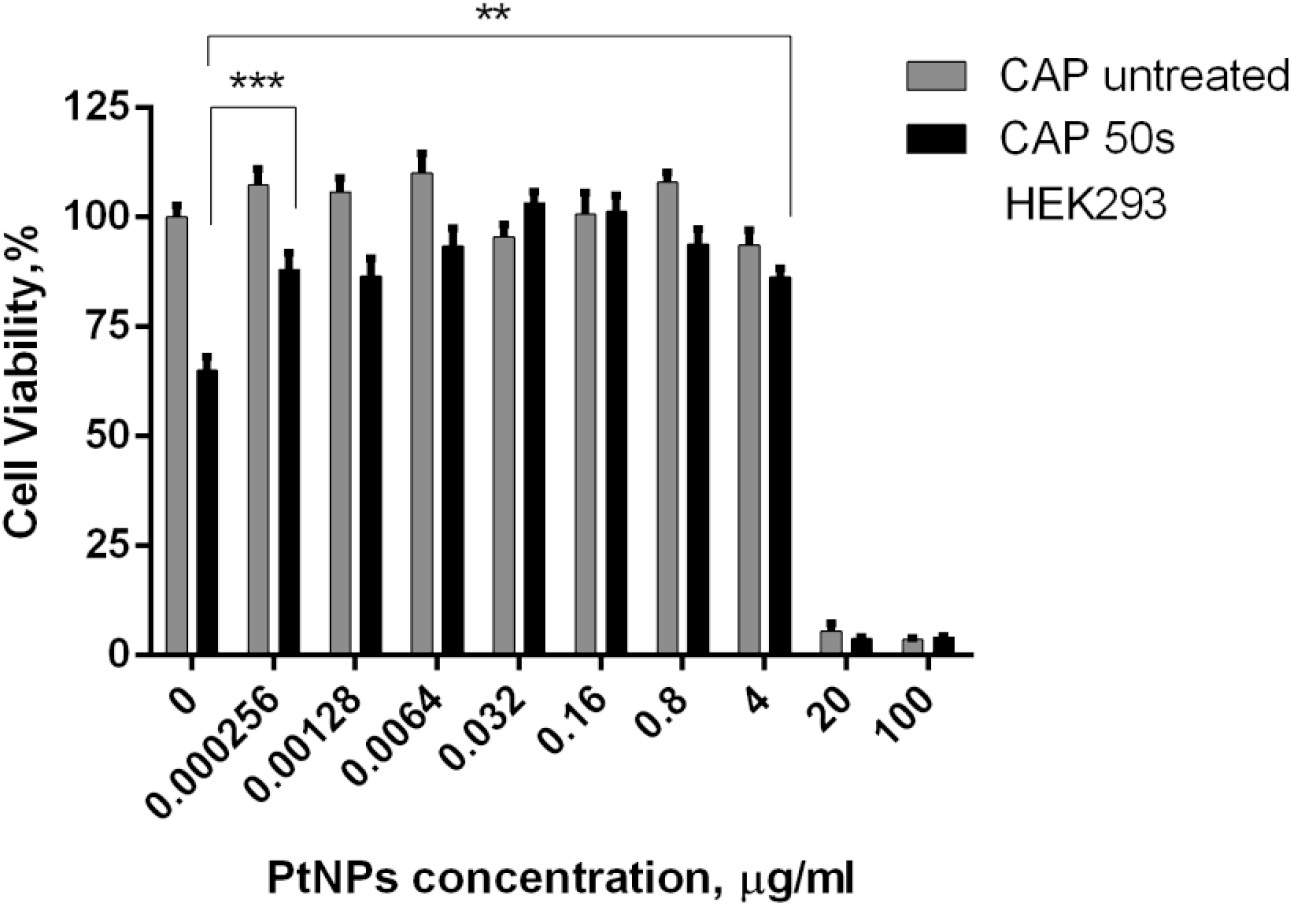
Dose response curves for PtNPs treatment in non-cancerous cells. HEK293 cells were incubated with corresponding amounts of PtNPs for 24h before CAP treatment and Alamar blue analysis was carried out 24h after CAP treatment. Black bars represent the combination of CAP treatment and PtNPs. Grey bars represent no CAP treatment. The first bar depicts negative control and the second black bar shows CAP treatment only with no 0 μg/ml PtNPs.

We observed a similar protective effect when HEK293 cells were loaded with PtNP before exposure to CAP for the concentrations of PtNP ranged from 4 μg/ml to 0.256 ng/ml (**P < 0.01, ***P < 0.001) compared with CAP only. The IC_50_ value of PtNP alone is 8.782 μg/ml (95% confident intervals = 6.529 μg/ml to 11.81 μg/ml) and the IC_50_ value of PtNP in combination with 50 s CAP treatment is 7.235 μg/ml (95% confident intervals = 5.366 μg/ml to 9.756 μg/ml) (Supplementary Figure S1b).

## Discussion

We have tested the PtNPs (3 nm) both with U-251 MG cancer cell line (Figure 1A, B) and HEK293 non-cancerous cell line (Figure 5) in combination with 50s CAP treatment. When the concentration of PtNP increased to 20 μg/ml and higher doses, cell viability rapidly decreased for both cell lines. We selected a dose of CAP close to the IC_50_ value in both cell lines, and as can be shown in figure 1A and figure 5, cytotoxicity measured was 50% and 40% for U-251 MG and HEK293 cells, respectively. Concentrations of PtNP at and below 4 μg/ml did not lead to significant cytotoxicity for both cell lines and our results are broadly consistent with other studies that indicate a low toxicity for PtNP^29,30,38^. We have observed similar IC_50_ values of PtNP alone and PtNP in combination with CAP treatment (Supplementary Figure S1a). Previous studies suggested that the size, shape, coating and surface area of noble-metal nanocrystals affect their catalytic properties^39^. For instance, Horie *et al*. (2011) treated A549 and HaCat cells with 5-10 nm pure PtNPs up to 17.4 mg/mL and did not see any cytotoxic effects, oxidative stress or cell death^29^. Pedone *et al*. (2017) showed that 5 and 20 nm citrate-capped PtNPs up to 100 μg/ml was not harmful to HUVEC cells^38^. Manikadan *et al*. (2013) characterised size-dependent toxicity using a range of PtNPs sizes, stabilized with PVP from 1 to 21 nm in Neuro 2 cells for 12 hours and showed that all of the sizes resulted in cellular damage except for 5-6 nm (up to 50 μg/ 1 × 10^6^ cells)^40^. Additionally, PVP-capped 5.8 nm PtNPs induced genotoxic effects and showed strong DNA damage at a concentration of 25 μg/ml but did not affect the cell viability at the time point of 24 h after PtNPs treatment in normal human epidermal keratinocytes^41^. Moreover, it was demonstrated that 1-5 nm PtNPs at concentrations up to 50 μg/ml did not show cytotoxic effects in TIG-1, HeLa, HepG2, WI-38 and MRC-5 cells^30^. Studies demonstrate that the size of PtNPs is one of the main factors on insignificant cytotoxicity even at higher concentrations^42,43^. Some *in vivo* studies demonstrated that small size PtNPs (2-19 nm) did not cause any side effects on chicken embryogenesis and up to 20 μg/ml concentrations did not alter liver functionality^44^. 3-10 nm PVP-PtNPs at concentrations up to 100 μg/ml did not show toxic effects on growth and development of zebrafish^45^. 1 nm PtNPs treated mice did not show any toxicity profile in their heart, lungs, spleen and liver however they suffered nephrotoxicity. Additionally, *in vitro* data reported that 8 nm PtNPs showed great biocompatibility^46^. Although many studies focus on the antioxidant effect of PtNPs *in vivo*, it has been reported that 21 nm PtNPs can induce inflammatory responses^47^. We purchased 3 nm commercial PtNPs with fully characterized size, shape, coating, purity, concentration to aid our toxicological profiling and allow for standardization of results.

CAP generates oxygen-based species (.OH), (^1^O_2_), (O_2_.^-^), (H_2_O_2_), (O_3_)^48^, as well as nitrogen-based species (NO), (NO_2_), (NO_3_), (N_2_O), (N_2_O_4_)^48^ in the gas phase. Physical elements include ultraviolet, heat, and electromagnetic fields in CAP. In addition to chemical and physical factors, positive charged ions such as N_2_^+49^ and electrons^50^ are also generated by CAP. The interaction of CAP-generated RONS and cells is an underlying principle for the anti-cancer effects of CAP^51^. Recently, many studies have focused on the potential selective killing effect of CAP in various cancer cells^50,52^. PtNP have previously been used for their antioxidant properties and it is proposed that PtNPs have a huge potential use as antioxidant nanodrugs^53^ by scavenging ROS in hepatic Kupffer cells^54^ or inhibiting apoptosis in human lymphoma U937 and HH cells^55^. Zheng *et al*. (2014) demonstrated that 2-4 nm PtNPs show synergistic scavenging activity with small antioxidant molecules^53^. PtNPs at low doses are capable of reducing ROS production caused by glucose, angiotensin and cholesterol treatments in endothelial cells^53^. Interestingly, Jawaid *et al*. (2016) reported that PtNPs show antagonism with Helium based CAP treatment. They demonstrated that PtNPs induced He-CAP desensitization in human lymphoma U937 cells and may inhibit pathways involved in apoptosis execution^56^. This study highlighted a very useful property of PtNP as possible catalase / SOD mimetics and their protective effect against CAP-induced cytotoxicity. It has been reported that PtNPs can catalyse the reduction of H_2_O_2_ to H_2_O and O_2_ by mimicking natural CAT. PtNPs are able to act as glutathione peroxidase, which launches the oxidation of a reduced substrate to decompose H_2_O_2_ to water, and they also catalyse the dismutation of O_2_ ^-^ into O_2_ and H_2_O_2_ as biological SOD mimics^35,57^. The potential of radical quenching ability of PtNPs, acting as enzyme mimetics, makes them promising candidates in cancer therapy and oxidative stress related diseases. However, it is not yet known whether this effect is observed in other CAP systems, or against adherent and non-cancerous cell lines.

We have investigated the potential antioxidant activity of PtNPs against ROS generated by DBD CAP system in U-251 MG and HEK293 cell lines. As Figure 2A presents, we detected significantly oxidised H_2_DCFDA in the presence of 50 s CAP treatment. Previous studies show that the combination of CAP and nanoparticles together proposes a strong anticancer capacity by showing synergistic effects^58^. However, as Figure 2B demonstrates, PtNPs dramatically reduced the CAP-induced ROS production in cells with three different concentrations of PtNPs (0.0512 ng/ml, 0.032 μg/ml, and 5 μg/ml) at concentrations that did not show any toxic effects in cells without CAP treatment. In addition, CAP-generated ROS levels were significantly decreased at a concentration of 0.032 μg/ml of PtNPs.

We found that the fluorescence level of H_2_DCFDA of CAP treated medium is not decreased by PtNPs in both increasing and/or decreasing (Figure 2C) concentrations of PtNPs, indicating the antioxidant effect of PtNP requires the nanoparticles either to be inside or near cells when exposed to CAP. One possibility is the mimetic activity of PtNP in cells. it has been reported that PtNPs act as a potent catalase mimetic at pH 7.4 and a potent peroxidase mimetic at pH 4.5^59^. The pH value of culture medium is usually 7.4, matching the pH of the cytosol and early endosomes. Other organelles such as late endosomes and lysosomes have much lower pH values, approaching 4.5 for lysosomes. Therefore, PtNP may act as either catalase or peroxidase mimetics in cells depending on their subcellular location. The localized accumulation of PtNPs in cellular compartments may drive localized antioxidant mimetic functions. Alternatively, PtNP may require the redox potential of living cells to fully catalyse redox reactions and quench reactive species^59^.

The PI staining measurements in Figure 3 show comparison of two different concentrations of PtNPs. PtNPs were not found to reduce the level of PI staining when compared with cells exposed only to 50 s CAP treatment. Our findings differ from Jawaid *et al*. (2016), who demonstrated using a He-CAP device that the PI staining was decreased in cells incubated with PtNPs during and after the CAP treatment, thus concluding that the fraction of secondary necrosis induced by 4 min He-CAP treatment was significantly decreased in the 300 μM PtNP treated group compared to non-PtNPs group^56^. This difference may be due to the differences in experimental design and methods. Jawaid *et al*. (2016) synthesised PtNP using citrate reduction but did not characterise the size of PtNPs, incubated cells with PtNP immediately after CAP treatment, and used different cell lines and plasma devices to those used in the current study^56^. As is seen in Figure 3B, C, we have analysed the percentage of dead and viable cells following PI staining and report that preincubation of PtNPs do not reduce the percentage of dead cells (no significant difference when compared to CAP treatment). However, as is seen in Figure 1, 4 and 5 PtNPs were found to protect total cell viability against CAP induced cytotoxicity and protect mitochondria membrane potential against CAP-induced oxidative stress. Therefore, it may demonstrate that PtNPs cannot reduce the percentage of dead cells as a result of CAP treatment but can protect the remaining viable cells and may ease the oxidative stress induced by CAP treatment.

We have previously demonstrated that 60 s CAP treatment induced rapid cell membrane damage, which was still evident up to 120 hours after exposure, corresponding to a loss of cell viability^36^. In contrast, a lower doses of CAP (30 s) induced membrane oxidation and triggered plasma membrane recycling and repair, but did not cause any overall increase in membrane permeability or loss of cell viability^12^. The inference was that the membrane damage observed was a causative factor in the loss of cell viability. We have demonstrated here that CAP treatment induces cell membrane damage in U-251 MG cells. Interestingly, CAP-induced membrane permeabilisation was not reduced by PtNPs, but other downstream indicators of viability such as JC-1 and Alamar blue were improved. It is likely that the PtNPs were mainly accumulated inside cells and therefore did not protect against initial membrane damage but prevented downstream ROS-dependent cytotoxicity.

We have previously demonstrated that the cell membrane peroxidation is induced by CAP-generated extracellular ROS^12^. In the current study, we demonstrate that PtNPs in culture medium possess low antioxidant ability and do not protect against membrane permeabilisation induced by CAP treatment. The pre-incubation of PtNPs with U-251 MG cells allowed the accumulation of PtNPs in cells, which leads to a more potent antioxidant response in cells compared to the culture medium treatment only. This observation along with the decreased intracellular ROS observed in CAP-treated cells pre-incubated with PtNP led us to investigate the role of mitochondria in the protection against CAP-induced cytotoxicity.

CAP-induced ROS can cause oxidative stress, which targets mitochondria and results in mitochondrial dysfunction and cell death^60^, and is presented as the loss of the mitochondrial membrane potential (ΔΨm)^37^. In agreement with these studies, we herein identified that CAP results in loss of mitochondrial membrane potential 24 h after the treatment in comparison with their untreated counterparts (Figure 4A and 4B) protected by low concentrations (0.032 μg/ml) of PtNP. Interestingly, high concentrations (5 μg/ml) of PtNP did not have a strong protective effect suggesting a possible disruption of normal cell function and viability occurs when using this higher dose of PtNP in conjunction with CAP.

In conclusion, we have demonstrated that, with intracellular accumulation, low doses of PtNPs showed non-to-low toxicity against U-251 MG and HEK293 cell lines while displaying significant protective effects against CAP induced cytotoxicity as a potent intracellular scavenger of ROS. With further investigation, PtNPs can be promising a candidate for cancer treatment by reducing the side effects of CAP therapy and other therapies that produce ROS by lowering intracellular ROS, preventing downstream cytotoxic signaling and providing cells time to repair the cellular damage caused by CAP.

## Acknowledgements

This work is supported by TU DUBLIN Fiosraigh Research Scholarship programme (S.G., Z.H.), Science Foundation Ireland Grant Number 14/IA/2626 (P.C., J.C.).

## Author Contributions

S.G., Z.H., R.M., P.C. and J.C. conceived the project and designed the experiments. S.G. and Z.H. performed the experiments, collected and analysed the data. S.G., Z.H., R.M., P.C., and J.C. co-wrote the paper. All authors discussed the results and reviewed the manuscript.

## Competing Interests

The authors declare no competing interests.

## Notes

### Competing Interest Statement

The authors have declared no competing interest.

