## Supplementary Material for "Platinum nanoparticles inhibit intracellular ROS generation and protect against Cold Atmospheric Plasma-induced cytotoxicity"

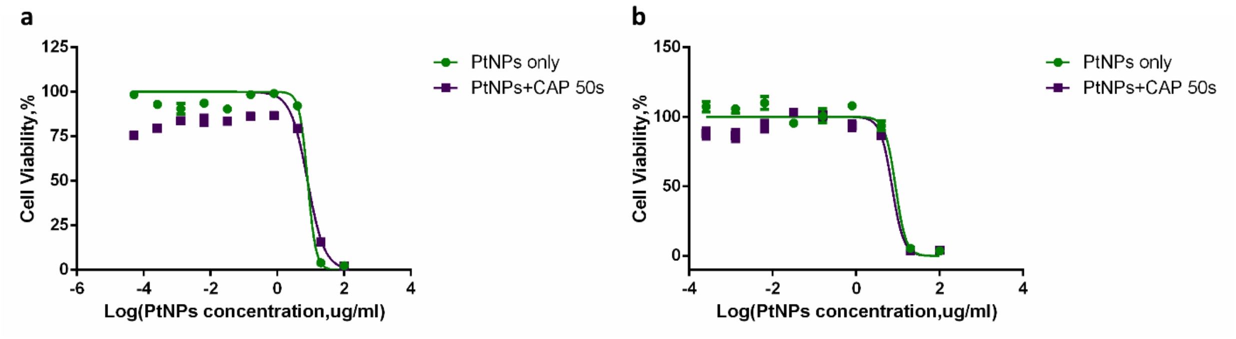


Supplementary Figure S1**.** Dose response curves for PtNPs treatment. U-251 MG cells (**a**) and HEK293 cells (**b**) cells were incubated with decreasing amount of PtNPs (starting from 100 µg/ml, 1:5 dilution) for 24h before CAP treatment. 24h after CAP treatment, Alamar blue analysis was carried out. All experiments were repeated at least three times. Data displayed was normalised to the untreated control and is shown as the % mean ± S.E.M.
